# SARS-CoV-2 infection activates endogenous retroviruses of the LTR69 subfamily

**DOI:** 10.1101/2023.03.21.533610

**Authors:** Ankit Arora, Jan Eric Kolberg, Smitha Srinivasachar Badarinarayan, Daksha Munot, Martin Müller, Daniel Sauter, Vikas Bansal

**Author notes:** Joint first authors (Ankit Arora, Jan Eric Kolberg).

## Abstract

Accumulating evidence suggests that endogenous retroviruses (ERVs) play an important role in the host response to infection and the development of disease. By combining RNA- and ChIP-sequencing analyses with RT-qPCR, we show that SARS-CoV-2 infection induces the LTR69 subfamily of ERVs, both *in vitro* and *in vivo*. Using functional assays, we identified one SARS-CoV-2-activated LTR69 locus, termed Dup69, which exhibits enhancer activity and is responsive to the transcription factors IRF3 and p65/RelA. LTR69-Dup69 is located about 500 bp upstream of a long non-coding RNA gene (ENSG00000289418) and within the *PTPRN2* gene encoding a diabetes-associated autoantigen. Both ENSG00000289418 and *PTPRN2* showed a significant increase in expression upon SARS-CoV-2 infection. Thus, our study sheds light on the interplay of exogenous with endogenous viruses and helps to understand how ERVs regulate gene expression during infection.

## Background

Severe acute respiratory syndrome coronavirus 2 (SARS-CoV-2), the causative agent of the COVID-19 pandemic, has caused unprecedented global health and socioeconomic impacts. With billions of infections and millions of reported deaths worldwide, there is a pressing need to better understand the complex interplay of SARS-CoV-2 with infected host cells and the pathogenesis of the disease.

Recent studies have suggested that repetitive DNA sequences known as transposable elements (TEs) play an essential role in the host response to viral infection and the development of disease. For instance, some TEs are capable of regulating the expression of antiviral factors and other host proteins through their activity as enhancers or promoters [1,2]. Furthermore, TE-derived nucleic acids may be sensed by cellular pattern recognition receptors and thereby amplify innate sensing cascades and the induction of Interferon-mediated immune responses [3]. In line with a potential role in the outcome of viral infections, viruses such as the Human Immunodeficiency Virus (HIV), Human Cytomegalovirus (HCMV) or Influenza A Virus (IAV) trigger the activation of transposable elements that are otherwise silenced [2,4–6].

Here, we leverage publicly available transcriptome and chromatin datasets of infected cell lines and patient-derived samples to decipher the impact of SARS-CoV-2 on the TE expression profiles of virus-infected or -exposed cells. Several studies have reported an induction of HERV-K [7–9], HERV-W [10,11] or HERV-L [12–16] upon SARS-CoV-2 infection. In line with this, we found that SARS-CoV-2 infection induces the activation of a particular subset of endogenous retroviruses (ERVs), so-called LTR69 repeats. These long terminal repeats (LTRs) represent solo-LTRs of the HERV-L family of endogenous retroviruses. In contrast to previous studies, we also performed mechanistic analyses and identified a SARS-CoV-2-responsive LTR69 repeat that acts as an enhancer and is activated by IRF3 and p65/RelA, two transcription factors that are activated upon sensing of viral RNA.

## Results and Discussion

To determine the effect of SARS-CoV-2 infection on the activity of TEs, we analyzed publicly available poly(A)-enriched mRNA-seq data obtained from cell lines and COVID-19 patients (Table S1). First, we took advantage of a data set from SARS-CoV-2-infected and uninfected Calu-3 cells to identify differentially expressed TEs. Calu-3 cells are a human lung cell line that is susceptible to SARS-CoV-2 infection *in vitro* and represents the natural target cells of the virus. Using TEtranscript, we found that solo-LTRs of two human endogenous retrovirus (HERV) subfamilies, LTR69 (Log_2_ FC = 3.11 and adjusted P-value = 1.06e-10) and LTR103_Mam (Log_2_ FC = 3.50 and adjusted P-value = 4.36e-27), were significantly up-regulated upon SARS-CoV-2 infection (Figure 1A and Table S2). Similarly, LTR69 expression was also elevated (Log_2_ FC > 1) in SARS-CoV-2 infected A549-ACE2 lung cells and bronchoalveolar lavage fluid (BALF) of deceased and non-deceased COVID-19 patients. In contrast, we observed no significant increase in LTR69 expression in Calu-3 cells that were infected with SARS-CoV or in peripheral blood mononuclear cells (PBMCs) of non-deceased COVID-19 patients, i.e. cells that are not susceptible to infection (Figure 1B). Since LTR69 repeats represent solo-LTRs of ERV-L, these results are in line with previous studies [12–16], which showed an up-regulation of ERV-L members upon SARS-CoV-2 infection. LTR103_Mam was significantly up-regulated in SARS-CoV-infected Calu-3 cells but not in BALF of non-deceased COVID-19 patients (Figure 1C). Using quantitative PCR (RT-qPCR), we validated the effect of LTR69 expression upon SARS-CoV-2 infection in both A549-ACE2 and Calu-3 cells (Figure 1D). While we observed only a modest elevation in Calu-3 cells, SARS-CoV-2 infection enhanced the LTR69 transcription about 4-fold in A549-ACE2 cells (P-value = 0.023604, one-sided t-test).

**Figure 1:**
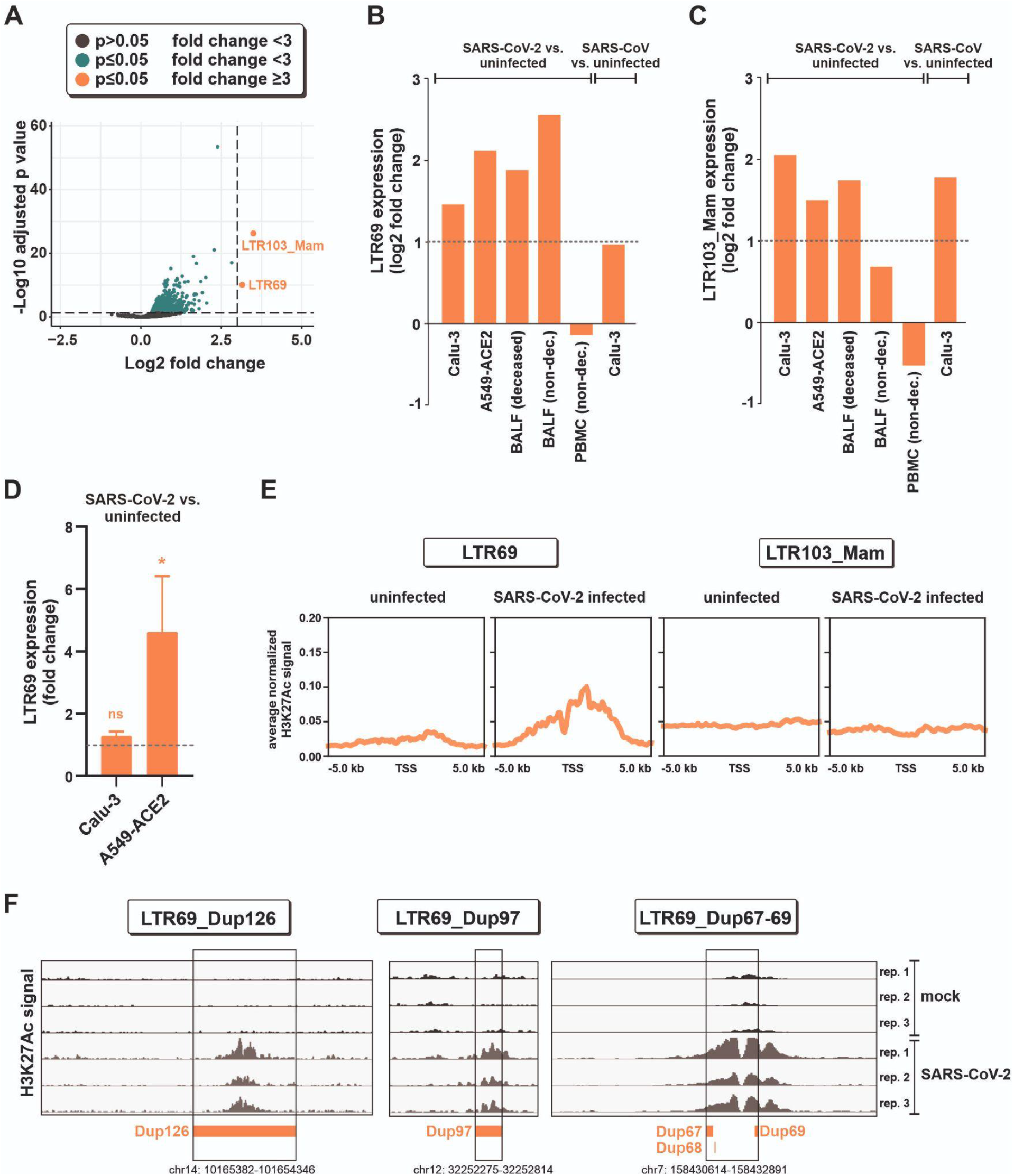
Activation of LTR69 repeats upon SARS-CoV-2 infection. **(A)** Volcano plot illustrating differential expression of transposable elements (TEs) in SARS-CoV-2-infected vs. uninfected Calu-3 cells (GSE147507). Dashed lines represent cutoffs of a log_2_ fold change of 3 and an adjusted P value of 0.05. **(B, C)** Log_2_ fold change of (B) LTR69 and (C) LTR103_Mam expression upon SARS-CoV-2 or SARS-CoV infection as determined in additional RNA-seq data sets (Table S1). Data sets were obtained from *in vitro* infected lung cell lines (Calu-3 cells (GSE148729), A549-ACE2 cells) or from bronchoalveolar lavage fluid (BALF) and peripheral blood mononuclear cells (PBMCs) *in vivo*. **(D)** Validation of SARS-CoV-2-induced LTR69 expression via qPCR in Calu-3 and A549-ACE2 cell. Cells were infected with an MOI of 2 and harvested 24 h post infection. Mean values of three independent experiments ± SEM are shown (* p<0.05, ns not significant). **(E)** An average H3K27ac signal profiling of LTR69 loci (left panel) and LTR103_Mam loci (right panel) around the transcription start site (TSS) is shown. ChIP-seq data were obtained in SARS-CoV-2 infected (24h, MOI 0.5) and uninfected A549-ACE2 cells. **(F)** Integrative Genomics Viewer (IGV) snapshots of exemplary H3K27ac peaks on individual LTR69 loci (hg38) in A549-ACE2 cells.

Notably, TEs can also be active without being transcribed. For example, many TE-derived elements act as enhancers regulating cellular gene expression [1]. One well-recognized epigenetic mark of active enhancers is the acetylation of lysine 27 in histone H3 (H3K27Ac). To dissect the enhancer activity of LTR69 and LTR103 in the presence and absence of SARS-CoV-2, we therefore analyzed publicly available chromatin immunoprecipitation sequencing data (ChIP-seq) of H3K27Ac in A549-ACE2 cells. Interestingly, the transcription start site (TSS) profile plot across all LTR69 loci (n=147) revealed an enrichment of H3K27Ac marks upon SARS-CoV-2 infection in comparison to uninfected cells (Figure 1E). In contrast, we did not observe any significant enrichment of enhancer marks across the LTR103_Mam loci (n=1262). Therefore, we focused our further analyses on individual LTR69 locus that showed at least one significant H3K27Ac peak identified by the MACS2 peak calling algorithm. There were 12 unique peaks of H3K27Ac on 15 LTR69 loci upon SARS-CoV-2 infection (Table S3). Exemplary peak signals are displayed as Integrative Genomics Viewer (IGV) screenshots in Figure 1F.

To test whether some of the SARS-CoV-2-activated LTR69 repeats exert regulatory effects, we tested them for potential enhancer activities. We selected five representative candidates (loci names defined in the annotation file as Dup60, Dup67, Dup68, Dup69, Dup97) (Table S3) and inserted them into enhancer reporter vectors. These plasmids express a *Gaussia* luciferase reporter gene under the control of a minimal promoter, whose activity may be increased by upstream enhancer elements (Figure 2A). Dup67 and Dup68 were inserted together into the same vector as they are located in close proximity in the genome (Figure 1F). A previously characterized LTR12C enhancer located upstream of the *GBP2* gene served as positive control [2]. As expected, LTR12C_*GBP2* increased *Gaussia* luciferase expression compared to the vector control lacking an LTR repeat (Figure 2B). A similar enhancing effect was observed for Dup69, whereas the remaining LTR69 elements had no significant modulatory effect or even decreased reporter gene expression. Since LTR69-Dup69 exerts enhancer activity, we hypothesized that it might regulate the expression of adjacent genes. Inspection of the respective gene locus revealed that Dup69 is located in an intron of *PTPRN2*, about 500 nucleotides upstream of ENSG00000289418 (Figure 2C). While *PTPRN2* codes for a tyrosine phosphatase receptor that serves as a major autoantigen in type 1 diabetes [17], ENSG00000289418 encodes a long non-coding RNA. We, therefore, analyzed the expression of both genes in SARS-CoV-2 infected versus uninfected lung cells via RT-qPCR. Intriguingly, expression of the lncRNA increased about 3.6- and 25.2-fold in Calu-3 and A549-ACE2, cells respectively (Figure 2D). Furthermore, expression of *PTPRN2* increased on average 4.1-fold upon SARS-CoV-2 infection in A549-ACE2 cells, while *PTPRN2* mRNA was not detectable in Calu-3 cells (Figure 2E). Interestingly, Sharif-Askari and colleagues also found up-regulation of *PTPRN2* in the whole blood of COVID-19 patients [18]. Although a causal link remains to be demonstrated, it is tempting to speculate that changes in the expression of ENSG00000289418 and/or *PTPRN2* are mediated by the enhancer activity of the LTR69-Dup69 repeat. To elucidate the mechanisms that may underlie the activation of LTR69-Dup69 upon SARS-CoV-2 infection, we screened its nucleotide sequence for binding sites of transcription factors that are known to be activated in infected cells. Using JASPAR [19], we identified putative binding sites for NF-κB subunits (NFKB1, NFKB2, Rel), IRF3 and STAT1 (Figure 2F). Intriguingly, LTR69-mediated enhancement of reporter gene expression could be further boosted by p65/RelA and a constitutively active mutant of IRF3, but not STAT1 (Figure 2G). In line with an activation of IRF3 and NF-κB upon innate sensing, the synthetic dsRNA analog polyI:C also significantly increased the activity of LTR69_Dup69 (Figure 2G). Together, these findings demonstrate that SARS-CoV-2 infection results in the activation of an LTR69 repeat that is responsive to p65/RelA and IRF3, acts as an enhancer element and may potentially regulate the expression of adjacent genes.

**Figure 2:**
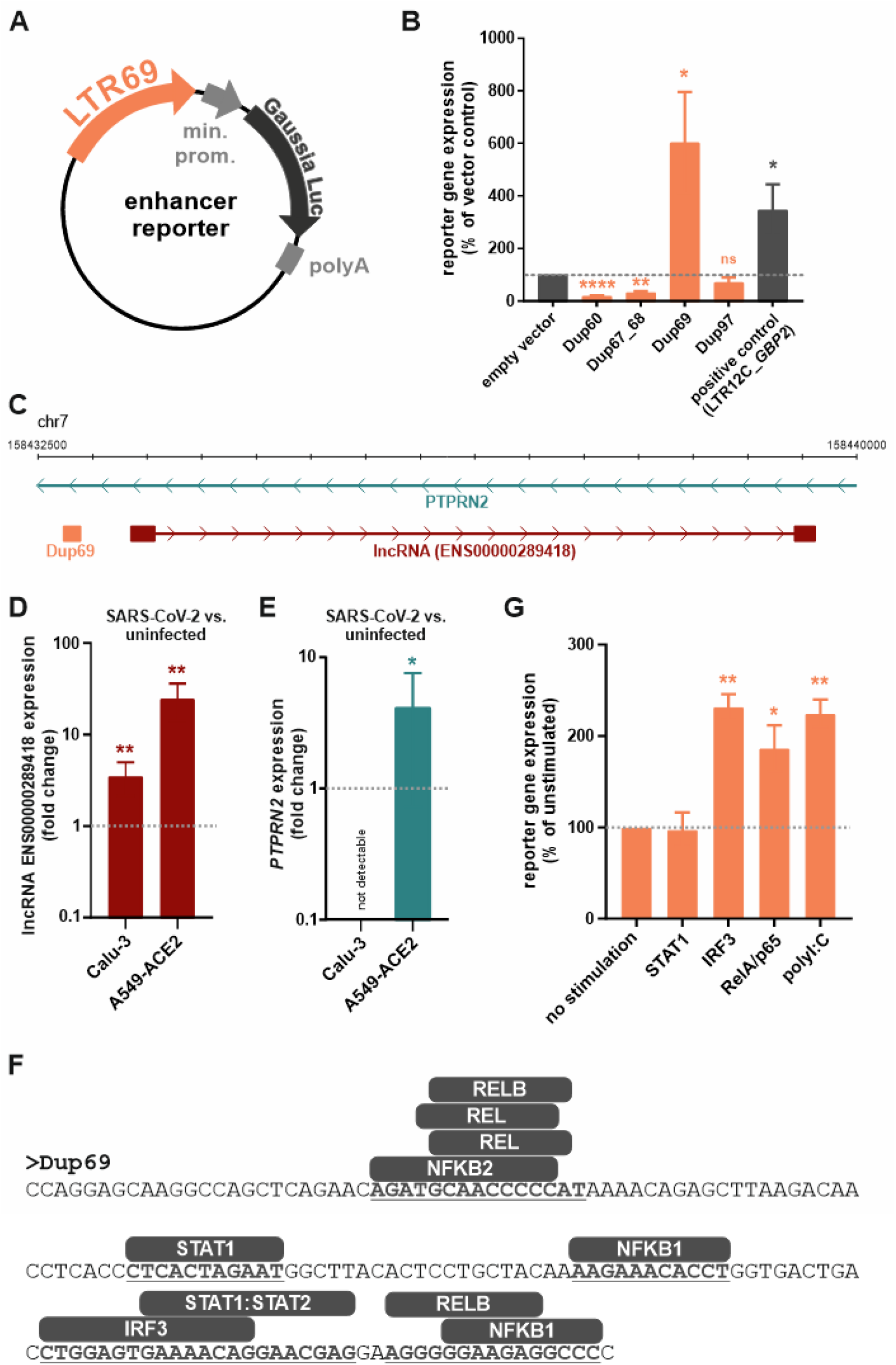
Enhancer activity of SARS-CoV-2-induced LTR69-Dup69. **(A)** LTR69 repeats (orange) were inserted into enhancer reporter vectors expressing *Gaussia* luciferase (black) under the control of a minimal promoter (grey). **(B)** HEK293T cells were co-transfected with the indicated reporter vectors expressing *Gaussia* luciferase and a control vector expressing firefly luciferase for normalization. A previously described LTR12C repeat with known enhancer activity served as positive control. Two days post transfection, reporter luciferase activity was determined and normalized to the activity of the control luciferase. Mean values of three to four independent experiments, each performed in triplicates are shown. Error bars indicate SD (* p<0.05, ** p<0.01, **** p<0.0001). **(C)** Integrative Genomics Viewer (IGV) snapshot (hg38) illustrating the localization of LTR69-Dup69 (orange) within an intron of the *PTPRN2* gene (green) and adjacent to the ENS00000289418 gene (red). **(D, E)** Expression of (D) ENS00000289418 and (E) *PTPRN2* in SARS-CoV-2-infected vs. uninfected Calu-3 and A549-ACE2 cells. Cells were infected with an MOI of 2 and harvested 24 h post infection. Mean values of two to three independent experiments ± SEM are shown. **(F)** Nucleotide sequence of LTR69-Dup69. The presence of putative binding sites for STAT1, NF-κB subunits and IRF3 is highlighted in bold and underlined. **(G)** HEK293T cells were co-transfected with the LTR69-Dup69 reporter vector expressing *Gaussia* luciferase, a control vector expressing firefly luciferase for normalization and increasing amounts of the indicated stimuli. Two days post transfection, reporter luciferase activity was determined and normalized to the activity of the control luciferase. Mean values of three independent experiments, each performed in triplicates are shown. Error bars indicate SD (* p<0.05, ** p<0.01, ns not significant).

Nevertheless, we would like to point out that this study is subject to limitations. Although we analyzed and compared multiple datasets, we acknowledge that the respective sample sizes are relatively small. In the future, it would be crucial to integrate and replicate the findings in other datasets, including cell-type specific data at both transcription and chromatin layers. We would also like to point out that expression of LTR69 and *PTPRN2* was elevated upon SARS-CoV-2 infection, but only reached relatively low mRNA levels. The extent of change in expression certainly depends on several factors such as the infected cell type, virus strain, multiplicity of infection (MOI) and time point post-infection. Moreover, computational analysis of TEs faces significant challenges due to a high false discovery rate [20]. While previous studies had already analyzed the effect of SARS-CoV-2 infection on TE activity, they have used different bioinformatics tools. For example, Marston and colleagues [15] used Telescope, which analyzes full-length transposable elements. Importantly, they also observed an induction of specific ERVs in SARS-CoV-2 infected or exposed cells, including ERV-L repeats.

Future studies will shed light on the downstream effects of TE activation on the virus and its host. It will be important to identify causal links between the activity of specific regulatory TEs (e.g. LTR69 repeats) and potential cellular target genes (e.g. *PTPRN2*). Furthermore, it will be interesting to investigate whether different viral infections trigger overlapping patterns of TE expression, indicative of a broader role of transposable elements in infection and immunity.

## Conclusions

In this short report, we confirm the differential expression and activation of specific mobile genetic elements in response to SARS-CoV-2 infection. In particular, we found and validated the up-regulation of the LTR69 subfamily of ERV-L upon infection. Moreover, we demonstrate that one of the SARS-CoV-2-induced LTR69 loci, LTR69-Dup69, exhibits enhancer activity and is responsive to the transcription factors p65/RelA and IRF3. LTR69-Dup69 is located about 500 bp upstream of a long non-coding RNA gene, ENSG00000289418, whose expression is also increased upon SARS-CoV-2 infection. At the same time, LTR69-Dup69 is located within an intron of the *PTPRN2* gene, which is also up-regulated upon SARS-CoV-2 infection and encodes for an autoantigen involved in type 1 diabetes. While further work is required, our study identifies LTR69 as a transposable element that is activated in SARS-CoV-2 infected cells and may modulate host gene expression and thus contribute to the outcome of SARS-CoV-2 infection.

## Methods

### Data Collection

The RNA-seq datasets analyzed in this study were downloaded either from GEO or the NCBI SRA database. Accession numbers and additional details are provided in Table S1.

### Transcriptome quantification and differential expression analysis

Raw RNA-seq reads from poly(A) RNA were subjected to a quality control using FastQC [v0.11.9] [21]. Quality controlled reads were subjected to removal of adapter sequences and quality filtering using fastp [v0.20.1] [22]. These filtered reads were mapped to the human (hg38) reference genome using bowtie2 [v2.3.0] [23]. Mapped files of technical replicates of any group were merged using samtools [v1.10] [24]. Differential expression analysis for transposable elements (TEs) were performed for mapped reads at the family level using TEtranscripts [v2.2.1] [25] in R [v3.3.0] with use of GTF downloaded from “https://hammelllab.labsites.cshl.edu/software/TEtranscripts” named as GRCh38_GENCODE_rmsk_TE.gtf.gz.

### ChIP-seq analysis

ChIP-seq data were downloaded from GSE167528 [26]. Raw ChIP-seq reads in fastq format were subjected to quality control using FastQC [v0.11.9]. Quality controlled reads were subjected to removal of adapter sequences and quality filtering using fastp [v0.20.1]. The filtered reads were mapped to the human (hg38) reference genome using bowtie2 [v2.3.0]. Peak calling on mapped reads was performed using MACS2 [v2.1.1] [27]. In addition, bigwig signals and matrix computation were performed for each condition using deepTools [v2.5.2] [28].

### Infection of Calu-3 and A549-ACE2 cells with SARS-CoV-2

One day before infection, A549-ACE2 (80,000 cells/well) or Calu-3 (200,000 cells/well) were seeded into a 48 well plate. Cells were infected with SARS-CoV-2 B.1 at an MOI=2 and incubated at 37°C for 24 h.

### RT-qPCR

RNA was isolated from infected cells 24 h p.i. using Qiagen RNeasy Mini Kit (Cat. #74106). gDNA was eliminated using Invitrogen DNA-*free*™ DNase Treatment & Removal (Cat. #AM1906). cDNA was synthesized using Applied Biosystems High-Capacity cDNA Reverse Transcription Kit with RNase Inhibitor (Cat. #4374966). qPCR was performed using New England Biolabs Luna® Universal qPCR Master Mix (Cat. #M3003L). All steps were performed according to the manufacturer’s instructions. The following primers were used:

LTR69 Fwd: 5’ GAA TTA CTG GGT CTC CAT GAC 3’
LTR69 Rev: 5’ AGG GAG TTT AAG CTA TTC TTG T 3’
ENSG00000289418 Fwd: 5’ GAA GTT TAC AGG CAA AAG CTG C 3’
ENSG00000289418 Rev: 5’ AAC CCA GTG CCA GGA ATG AA 3’
GAPDH Fwd: 5’ GAG TCC ACT GGC GTC TTC A 3’
GAPDH Rev: 5’ GGG GTG CTA AGC AGT TGG T 3’

The following primer-probes were used for the PTPRN2 qPCR:

PTPRN2 FAM ThermoFischerScientific (Cat. #4448892; AssayID: Hs00243067_m1)
GAPDH VIC ThermoFisherScientific (Cat. #4448489)

All reactions were performed in duplicates, and GAPDH was used to normalize RNA expression across all samples. LTR69 primers were designed and used previously by Mao and colleagues [29]. Raw RT-qPCR data is provided in Table S4.

### Reporter plasmids

To generate promoter reporter vectors, LTR69 and LTR12C loci were synthesized (GenScript) and inserted into the pGLuc Mini-TK 2 *Gaussia* luciferase enhancer reporter plasmid (NEB) via KpnI/SacI restriction sites, upstream of the minimal promoter, as previously described [2].

### Cell culture

Human embryonic kidney 293T (HEK293T) cells and A549-ACE2 cells were cultured in Dulbecco’s modified Eagle medium (DMEM) containing 10% heat-inactivated fetal calf serum (FCS), 2 mM glutamine, 100 μg/ml streptomycin and 100 units/ml penicillin. HEK293T cells were tested for mycoplasma contamination every three months. Only mycoplasma negative cells were used for this study. Calu-3 were cultured in Dulbecco’s modified Eagle medium (DMEM) containing 10% heat-inactivated fetal calf serum (FCS) plus 2 mM glutamine, 100 μg/ml streptomycin and 100 units/ml penicillin. Medium was changed daily.

### Prediction of transcription factor binding sites

Putative binding sites for NF-κB subunits, IRF3 and STAT1 were predicted using JASPAR 2022 [19]. *Homo sapiens* was selected as species, and the relative profile score threshold was set to 70%. The ten sequence motifs with the highest relative scores (0.74-0.80) are shown in Fig. 2F.

### Enhancer reporter assay

HEK293T cells were seeded (30,000 cells/well) in poly-L-lysine coated 96-well tissue culture plates. After 24 hours, cells were transfected with a combination of expression vectors expressing *Gaussia* luciferase under the control of a minimal herpes simplex virus (HSV) thymidine kinase promoter (25 ng), either alone or upstream of a LTR12C or LTR69 locus, as well as a pTAL firefly luciferase plasmid (50 ng) as a normalization control and polyI:C (2,000 ng) or an expression plasmid for p65 (100 ng), STAT1 (100 ng) or a constitutively active mutant of IRF3 (1,000 ng). After 24 h, supernatants were harvested, and cells were lysed in 100 μl 1x Passive Lysis Buffer (Promega). *Gaussia* luciferase activity in the supernatants was measured by addition of Coelenterazine (PJK Biotech) and firefly luciferase activity was measured in the cells using the Luciferase Assay System (Promega) according to the manufacturer’s instruction.

## Ethics approval and consent to participate

Not applicable.

## Consent for publication

Not applicable.

## Availability of data and materials

All data generated or analyzed during this study are included in this published article [and its supplementary information files].

## Authors’ contributions

A.A. and V.B. conceived the project. D.S. and V.B. supervised the study. A.A. performed all the computational analysis. J.E.K. performed most of the RT-qPCR experiments and some of the luciferase reporter assays. M.M. generated SARS-CoV-2 virus stocks and performed some of the infection experiments. S.S.B. cloned enhancer reporter plasmids and D.M. performed some of the luciferase reporter assays. D.S. provided resources for the experimental work. A.A., J.E.K., D.S. and V.B. wrote the manuscript. All authors contributed to and reviewed the manuscript.

## Acknowledgments

We thank Isabell Haußmann and Corinna Bay for technical support. A549-ACE2 and Calu-3 cells were kindly provided by Michael Schindler; the IRF3 and p65 expression plasmids were kindly provided by Konstantin Sparrer and Bernd Baumann, respectively. We also thank Armin Ensser for providing SARS-CoV-2 B.1.

